# Six1 Promotes Skeletal Muscle Thyroid Hormone Response through Regulation of the MCT10 Transporter

**DOI:** 10.1101/2021.08.27.457933

**Authors:** John Girgis, Dabo Yang, Imane Chakroun, Yubing Liu, Alexandre Blais

## Abstract

The Six1 transcription factor is implicated in controlling the development of several tissue types, notably skeletal muscle. Six1 also contributes to muscle metabolism and its activity is associated with the fast-twitch, glycolytic phenotype. Six1 regulates the expression of certain genes of the fast muscle program by directly stimulating their transcription or indirectly acting through a long non-coding RNA. Under the hypothesis that additional mechanisms of action might be at play, a combined analysis of gene expression profiling and genome-wide location analysis data was performed. The *Slc16a10* gene, encoding the thyroid hormone transmembrane transporter MCT10, was identified as a gene with a transcriptional enhancer directly bound by Six1 and requiring Six1 activity for full expression in adult mouse tibialis anterior, a predominantly fast-twitch muscle. Of the various thyroid hormone transporters, MCT10 mRNA was found to be the most abundant in skeletal muscle, and to have a stronger expression in fast-twitch compared to slow-twitch muscle groups. Loss-of-function of MCT10 in the tibialis anterior recapitulated the effect of Six1 on the expression of fast-twitch muscle genes and led to lower activity of a thyroid hormone receptor-dependent reporter gene. These results shed light on the molecular mechanisms controlling the tissue expression profile of MCT10 and identify modulation of the thyroid hormone signaling pathway as an additional mechanism by which Six1 influences skeletal muscle metabolism.

## Introduction

The Six family of transcription factors (TFs) are a group of six homeodomain proteins that share a conserved, characteristic Six domain (1,2). They were discovered through complementation experiments to the *Drosophila sine oculis* gene, which is essential for the development of the fruit fly compound eyes (3). Homologues of the Six family of proteins have been discovered across the animal kingdom (4–6) and strong evidence exists that the Six TF-related regulatory network developed evolutionarily prior to the divergence of vertebrates (7). Six family gene knockout models highlight that these TFs play crucial roles at a global level in the development of several tissue types including skeletal muscle (8), kidney (9,10), neurones (11), cardiac muscle (12), eyes (13), and gonads (14).

In skeletal muscle, *Six1* and *Six4* have been shown to be involved in the transcriptional expression of Myogenin (15–17), a myogenic regulatory factor (MRF) essential for terminal differentiation of skeletal myofibers (18,19). Cooperative interactions between the MRFs and the Six TFs regulate downstream targets of both families of TFs, expressing key genes in the processes of muscle stem cell proliferation, fusion, and differentiation (17,20–23). The role of *Six1* in developing muscle is essential, as *Six1* knockout mice die at birth due to improperly formed diaphragms (24). In adult muscle stem cells, Six1 and Six4 are both essential for tissue regeneration following an injury (22,25). Comparatively, the role of the Six TFs in adult muscle homeostasis is arguably less well understood. In adult mature skeletal muscle, forced expression of *Six1* and its cofactor *Eya1* is sufficient to reprogram slow-twitch oxidative muscles towards a fast-twitch glycolytic phenotype (26). Conversely, myofiber-specific *Six1* conditional knockout (Six1-cKO) mice exhibit a switch towards a slow-twitch muscle phenotype, typified by expression of slow-twitch Myosin and Troponin isoforms (27,28).

The main proposed mechanism for *Six1* regulation of adult muscle fiber-type is through direct and positive transcriptional regulation of Linc-MYH, a long non-coding RNA (28). Linc-MYH knockdown recapitulates a majority of *Six1* knockout condition phenotypes, albeit to about 80% of the magnitude of effect, and is incapable of significantly increasing slow-twitch Myosin isoform expression. This partial recapitulation of the *Six1* knockout phenotype suggests that additional pathways that regulate fiber-type specification may simultaneously be under *Six1* regulatory control. While many regulatory networks are implicated in fiber-type maintenance in adult skeletal muscle, including Calcineurin/NFAT signaling (29), the AMPK axis (30), PGC-1α (31), Sox6 (32), and thyroid hormones (TH) signaling (33), the exact hierarchy and interplay between the different signalling and transcription pathways in the maintenance of adult muscle phenotype is unclear.

Here, we report findings on the implication of *Six1* in regulating the TH pathway. Similar to the Six TFs, TH play a crucial role in mammalian health, participating in the development, differentiation, growth, and metabolic homeostasis of various tissues, including the brain, muscles, liver, and pancreas (34). TH exist in two main forms, T_4_ and T_3_ and can affect cellular metabolism either through genomic or nongenomic regulation (35). While T_4_ has been shown to have a larger role in nongenomic interactions (36), T_3_ is the main genomically active form of TH acting as a ligand to activate thyroid receptor TFs (coded by the *Thra* or *Thrb* genes) to bind TH response elements (TREs) in the regulatory regions of target genes and modulate their transcription levels (37).

In skeletal muscles, TH activity regulates the processes of myogenesis and muscle regeneration (38,39), contractile structure (40), energy expenditure (41), mitochondrial thermogenesis, fatty acid oxidation (42), glucose uptake (40), insulin responsiveness (43), and autophagy (44). In the context of muscle fiber type specification, hyperthyroidism promotes fast-twitch muscle phenotypes, while hypothyroidism is associated with a switch towards slow-twitch phenotype (45,46). Mechanistically, TH receptors have been shown to directly control the expression of certain genes associated to the fast-twitch phenotype (47) and to indirectly antagonize the slow-twitch program by upregulating miR-133a1, which in turn downregulates Tead1, a TF driving the slow-twitch program (33,48). T_3_ has been demonstrated to regulate the expression of multiple MRFs, supported by the discovery of functional TREs driving mRNA transcription of *Myod1* (49) and *Myog* (50). Hyperthyroidism has been shown to push satellite cells towards expressing higher amounts of *Myod1* (51), and conversely, hypothyroidism impedes myogenic differentiation (52).

Due to the inability of TH to passively diffuse across lipid bilayers, tissue-specific hormone transporters are required for the cellular efflux and uptake of both T4 and T3 (53). Several proteins have the ability to transport TH across membranes. They differ in their substrate specificity, tissue expression profile and whether they operate primarily in hormone influx or efflux (54). MCT8, MCT10, OATP1C1, LAT1 and LAT2 (encoded by the genes *Slc16a2, Slc16a10, Slco1c1, Slc7a5* and *Slc7a8*, respectively) are the most relevant to TH cellular influx (53). MCT8 knockout mice present with a regulatory shift away from slow-twitch towards fast-twitch skeletal muscle gene expression (55,56). Interestingly, MCT8 is expressed only at low levels in skeletal muscle, and the effect of MCT8 loss-of-function in this tissue are explained by the increased circulating T3 levels in these animals, driving hormone influx and downstream gene activation (55). MCT8-deficient mice also display impaired muscle regeneration following a myotrauma (57). MCT8 is detected in muscle satellite cells, the adult stem cells responsible for this process, and specific deletion of MCT8 in this cell type impairs their differentiation, indicating that the regeneration defects in constitutive MCT8 knockout mice are not exclusively due to their above-mentioned serum TH imbalance(57). MCT10 is a protein structurally related to MCT8 and is expressed at higher levels in skeletal muscle, compared to MCT8 (54). Yet, less is known about the role MCT10 plays in skeletal muscles. It has been demonstrated recently that in the gastrocnemius muscle, the mRNA levels of MCT10, but not those of MCT8, increase linearly during the aging process of mice, suggesting a potential homeostatic regulatory role for MCT10 (58). OATP1C1 expression in rodents is mostly restricted to the brain, but its expression has also been detected in muscle satellite cells that are activated by *in vitro* culture(57). Finally, LAT1 and LAT2 can also participate in TH uptake (59,60). While LAT2 is the most highly expressed of the two in skeletal muscle (53), the mRNA of LAT1 appears more abundant than that of LAT2 in muscle satellite cells (57). Muscle-specific knockout of LAT1 blunts mTOR-S6K pathway activation by leucine exposure in this tissue(61).

There is paucity of studies on the mechanisms underlying the tissue-specific expression profiles of the various TH transporters (54,62). Additionally, to our knowledge no study has directly investigated links between Six TF and TH functions. The overlapping regulatory roles of both pathways in the regulation of MRF expression (15,50) and skeletal muscle type specification (28,33,63,64), suggests an interplay between the two regulatory networks worth investigating. With this study, we demonstrate that the thyroid transporter MCT10 is under direct transcriptional control of Six1 in adult skeletal muscle.

## Results and Discussion

### Strong correlation between Six1 function and the fast-twitch muscle program

We were initially interested in identifying genes under the direct control of Six1 and which might be associated with the fast-twitch phenotype. For this, we analyzed two gene expression profiling datasets. First, the dataset of Sakakibara et al. provides microarray gene expression profiles in gastrocnemius muscle in wild-type and in muscle-specific Six1 knock-out adult mice (Six1-cKO)(28). Second, the muscleDB dataset from Terry et al. provides RNA-seq expression profiles in several different murine muscle groups (65). We started by identifying genes differentially expressed in Six1-cKO gastrocnemius and found 204 genes requiring Six1 for normal expression levels (Figure 1, left-hand side heatmap; 86 genes less expressed in Six1-cKO and 118 more expressed). To examine in what proportion the genes deregulated in Six1-cKO gastrocnemius are associated with fast- or slow-twitch specific expression programmes, we considered their expression in the muscleDB dataset. We limited the analysis to a subset of muscle groups that are predominantly fast-twitch (EDL, quadriceps, TA, gastrocnemius, plantaris, classified in cluster 1 in muscleDB) or slow-twitch (soleus, FDB, diaphragm, classified in cluster 2 in muscleDB). We observed a striking association between gene downregulation in the absence of Six1 and genes being normally more highly expressed in fast-twitch muscles; and vice-versa between upregulation in the Six1-cKO and higher gene expression in slow-twitch muscles (Fig. 1, right-hand side heatmap). A Pearson correlation coefficient of -0.72 with two-tailed T test p-value smaller than 0.05 was calculated between the log fold-changes (Six1-cKO minus WT and fast minus slow) of these genes (Fig. 1, histogram).

**Figure 1.**
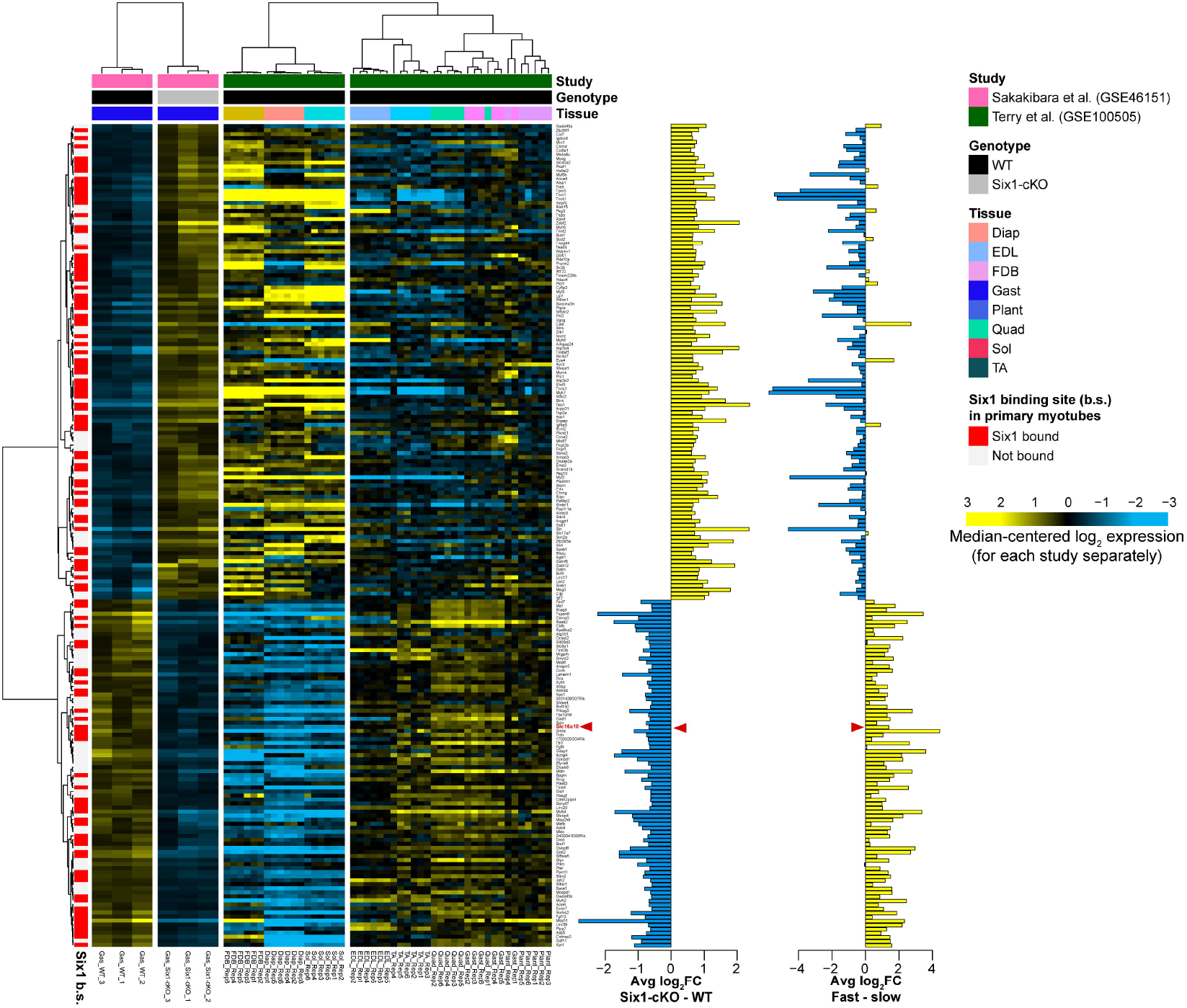
Skeletal muscle group distribution and Six1 dependence of gene expression. DNA microarray gene expression profiling from Sakakibara et al. were analyzed to identify differentially expressed genes in the gastrocnemius muscles of wild-type versus Six1-cKO mice. The cut-offs used were abs(log_2_FC) > 0.58 and Benjamini-Hochberg adjusted p value < 0.05. Genes were annotated based on whether their transcription start sites are within 50 kb of a Six1 binding site in primary myotubes (red bars). Pearson correlation hierarchical clustering was applied to rows (genes) and columns (samples). To further annotate these Six1-dependent genes, their expression from RNA-seq performed in several muscle tissue groups from the muscleDB database (Terry et al.) was also plotted. The rows retained the clustering solution order from the Sakakibara study, and columns were independently clustered by Pearson correlation (clustering performed separately from the Sakakibara data). The log_2_ fold-changes of each gene in the two experiments are shown in the vertical histograms, blue and yellow bars indicating negative and positive values, respectively. A red arrowhead indicates where the *Slc16a10* gene appears in these graphs. The gene symbols in this figure are also listed in supplementary table S3.

To increase the likelihood that the genes identified by this analysis are direct transcriptional targets of Six1, the genes were annotated for proximity to a Six1 binding site in ChIP-seq in primary mouse myotubes (Fig. 1, red bars in the row annotations). More than half of the differentially expressed genes (115 out of 204) have a Six1 binding site in their vicinity. Interestingly, genes upregulated in Six1-cKO were as likely to be bound by Six1 (71 of 118 genes have a Six1 binding site) as the genes downregulated in Six1-cKO (44 of 86).

These results not only confirm that *Six1* is implicated in establishing the proper gene expression profile of several genes in gastrocnemius muscle (28), but also reaffirm *Six1*’s function as both activator and repressor of gene expression. Previous reports have shown that *Six1* and other Six family TFs may repress transcription in cooperation with co-regulators *Sobp* (66) and members of the Dach or Groucho/TLE families (15,67), and activate transcription in association with co-factors of the Eya family or the SWI/SNF complex (15,68–71). This association between *Six1* binding and both gene up- and down-regulation after loss-of-function was previously observed in cultured myoblasts and myotubes (17) but the precise mechanisms underlying transcriptional repression by Six1 in skeletal muscle, and how they may be implicated in determining the slow- or fast-twitch expression programs, remain to be uncovered. A limitation of our study is that our *Six1* ChIP-seq analysis was performed in cultured primary cells differentiated *in vitro*, representing a heterogenous mixture of muscle precursors coming from different muscle groups. It is possible that our *Six1* genomic binding data reflects not only maintenance of a differentiated myofiber phenotype such as the fast-twitch program, but also the implication of *Six1* in upregulating these genes during the myogenic differentiation process. A genome-wide study of *Six1* binding performed in myofibers isolated from specific muscle groups will be required to further assess the function and mode of action of this TF in establishing and maintaining the slow-versus fast-twitch gene expression programmes.

### Six1 directly regulates expression of the MCT10 transporter in skeletal muscle

We focused our attention on identifying regulators of the fast-twitch programme that might be under direct control of Six1. Our attention was drawn to *Slc16a10* gene (marked with red arrowhead in Fig. 1), which became the focal point of the present study. *Slc16a10* encodes the MCT10 transmembrane transporter for TH, which are linked to fast-twitch skeletal muscle formation (34). Our analyses found that the expression of MCT10 is downregulated in Six1-cKO gastrocnemius muscle (Fig. 1). Among the five major genes encoding TH influx transporters, MCT10 is expressed in skeletal muscle to considerably higher levels than the other four tested (54). Further, analysis of muscleDB RNA-seq data showed that MCT10 expression is noticeably higher in the fast-twitch muscle groups compared to the slow ones (Fig. 1 and 2A). This fiber type specific profile was validated using quantitative reverse-transcription PCR (qRT-PCR) on RNA isolated from gastrocnemius, soleus and tibialis anterior of mice (Fig. 2B). These results, along with prior knowledge on the role of TH in muscle metabolism, suggest that MCT10 may be a mediator of Six1’s effect in establishing the fast-twitch phenotype.

**Figure 2.**
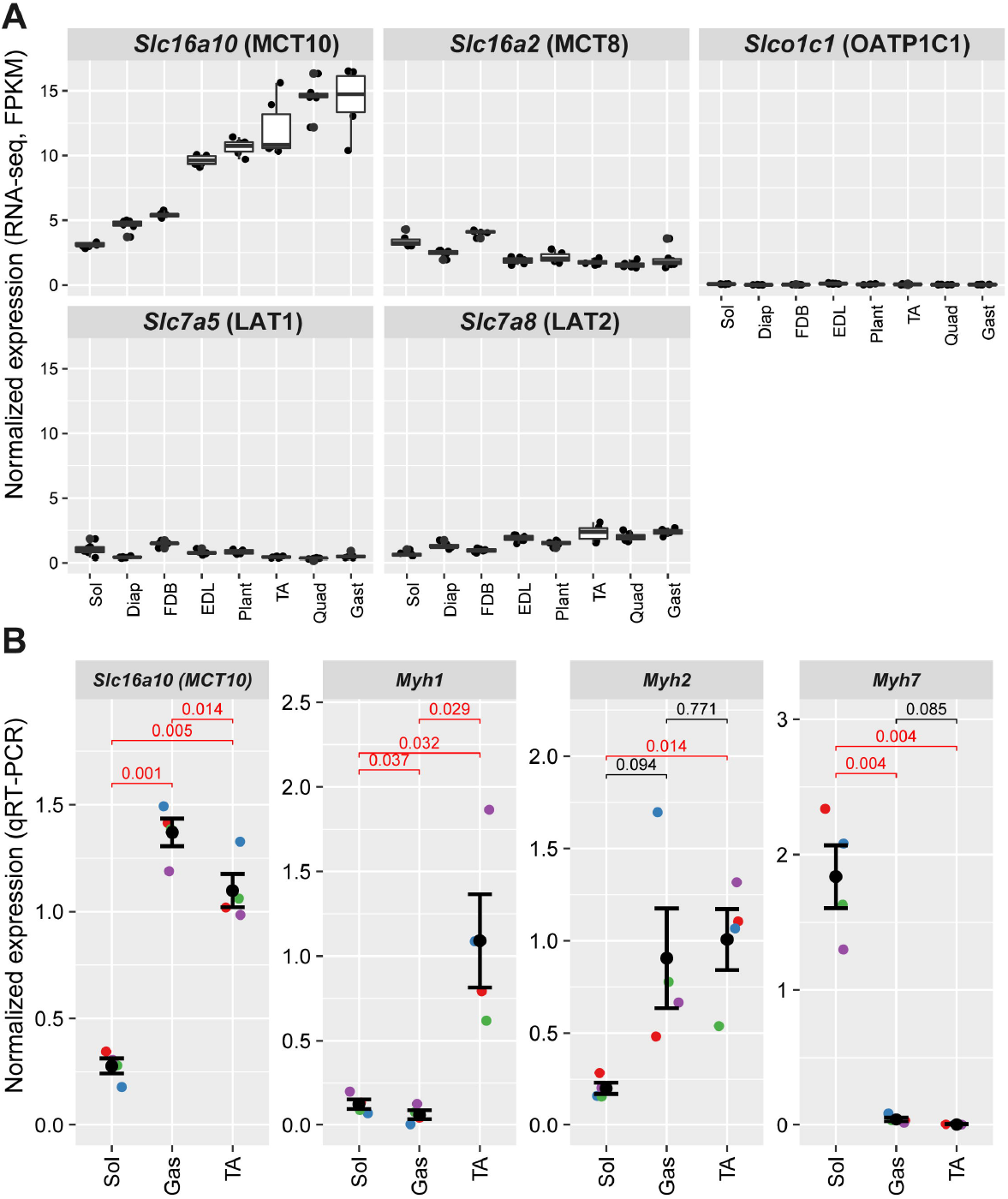
Expression of thyroid hormone transporter-coding genes in select murine skeletal muscle groups. **A)** RNA-seq expression data from muscleDB were quantitated and normalized by fragments per kilobase of transcript per million mapped reads (FPKM). The expression of the five transporters is shown: Slc16a10 (MCT10), Slc16a2 (MCT8), Slc7a5 (Lat1), Slc7a8 (Lat2) and Slco1c1 (Oatp1c1). Each of six replicates is represented by a dot and the statistical distribution of FPKM values is represented by the boxplot. **B)** qRT-PCR validation of *Slc16a10* expression in soleus (Sol), gastrocnemius (Gas) and tibialis anterior (TA) muscles. Expression was normalized over the geometric mean of control genes *Actnb* and 18S rRNA. Expression in soleus and tibialis anterior is lower than in gastrocnemius, by paired, two-tailed t test (p value < 0.05 shown in red). The expression profiles of *Myh1, Myh2* and *Myh7* is shown for comparison.

By examining our ChIP-seq data, we noted the presence of a *Six1* binding site located 48 kilobases upstream of the transcription start site of the MCT10 gene (Fig. 3). To obtain additional evidence that this *Six1* binding site represents a regulatory element, we analyzed histone mark ChIP-seq and DNA accessibility (ATAC-seq) data acquired in fast-twitch quadriceps and in slow-twitch soleus (72). The *Six1* binding site is located within a likely gene enhancer, as it is characterized by high DNA accessibility, the presence of H3K3me2 and H3K27ac, and these three marks are all more abundant in the quadriceps samples than the soleus samples (Fig. 3). To validate the *Six1* binding site identified by ChIP-seq, we performed a ChIP assay with anti-*Six1* and control rabbit IgG on chromatin isolated from pooled mouse hindlimb skeletal muscle tissue. This was followed by quantitative PCR on the ChIP eluates (qChIP) using primers specific for this -48kb enhancer, a positive control (the Myog promoter (17)) and a negative control locus (Hoxd10 locus, heterochromatic in skeletal muscle (73,74)). This showed a significant enrichment of *Six1* binding (Fig. 4). Our results confirm that the -48kb putative enhancer at the MCT10 locus is a *bona fide* Six1 binding site.

**Figure 3.**
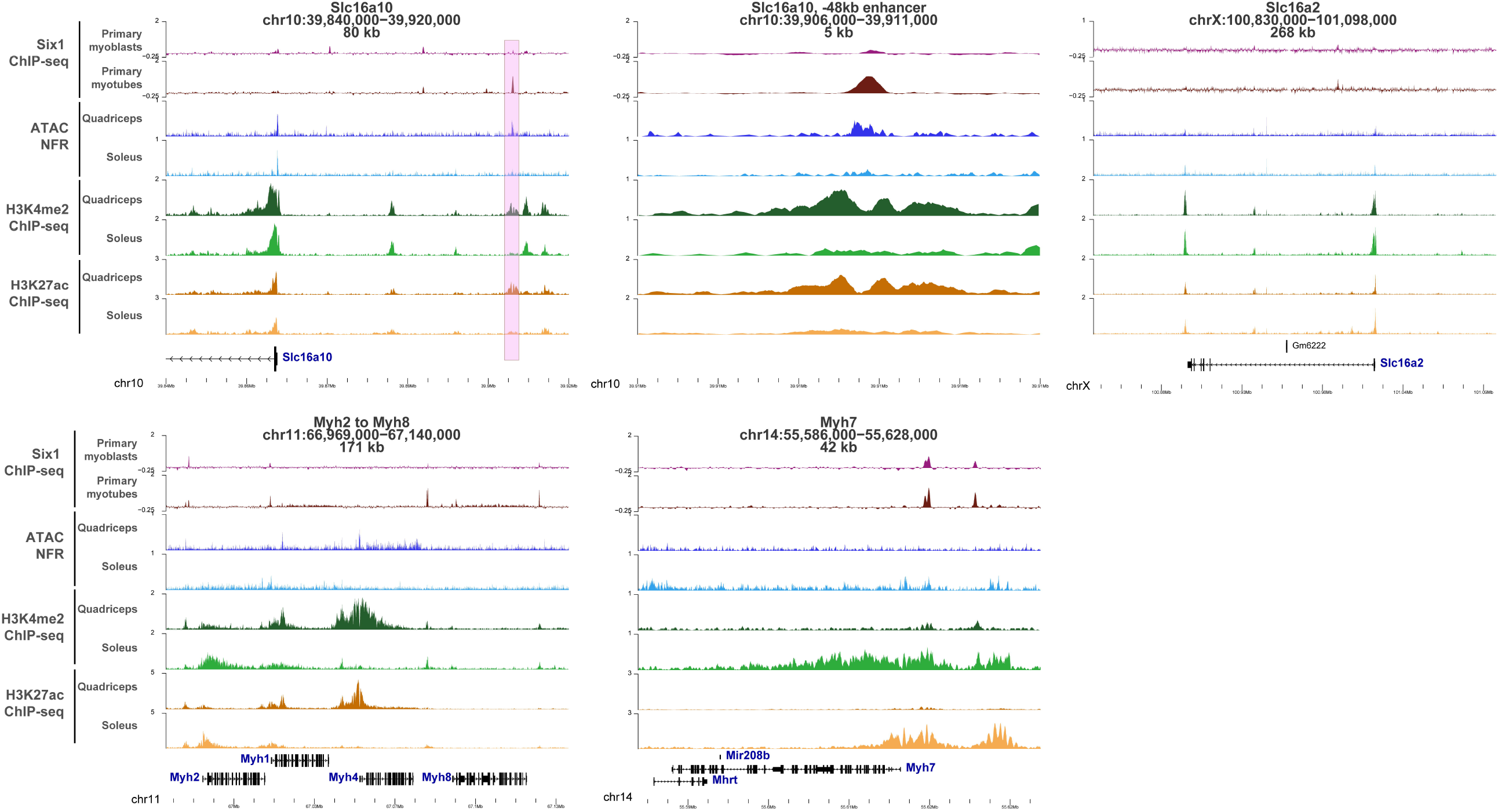
Six1 binding profile and epigenetic marks at the genes coding the two main thyroid hormone transporters. At each locus, the gene name, genomic position (mm9 coordinates) and length of the interval shown are given. The Six1 ChIP-seq signal represents data from primary proliferating myoblasts or differentiated myotubes, quantitated in sliding windows across the genome in reads per million sequenced, and subtracted from the signal in input chromatin control sample. The ATAC-seq data in quadriceps and soleus is shown for inferred nucleosome-free regions (insert sies between 10 and 130 base pairs) and quantitated in bins of 10 base pairs in counts per millions sequenced (CPM). The H3K4me2, indicative of enhancers and promoters, and H3K27ac, indicative of active promoters and enhancers, are quantitated in bins of 10 and CPM. The pink box upstream of the Slc16a10 start site shows the location of a putative Six1-bound enhancer, active predominantly in fast-twitch quadriceps, compared to slow-twitch soleus. This region is shown in detail in the plot to the right (“Slc16a10, -48kb enhancer”). The Myosin heavy chain loci are shown for comparison (bottom row).

**Figure 4.**
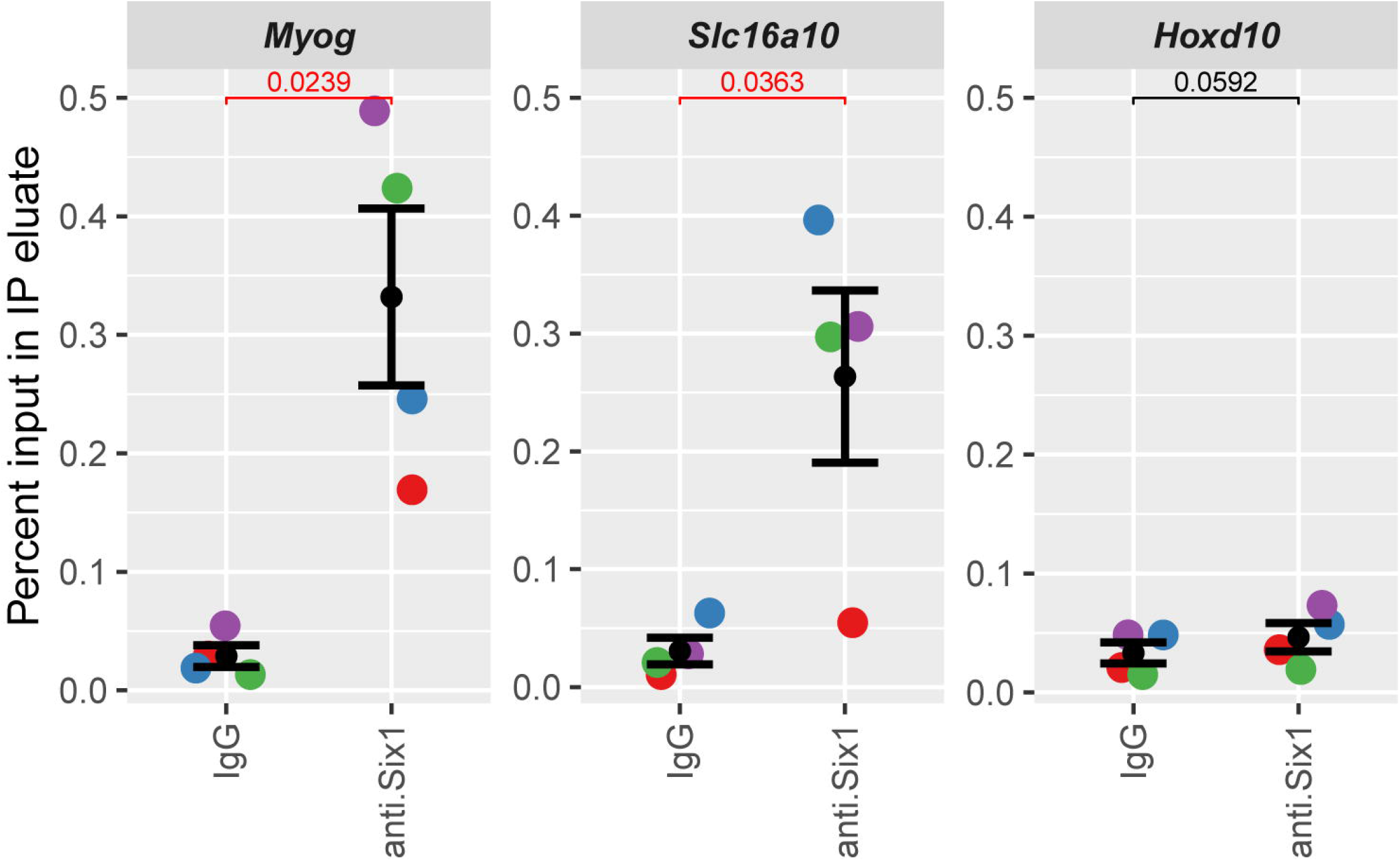
Confirmation of Six1 binding at the Slc16a10 enhancer. Chromatin from hindlimb muscles of four mice was harvested, fixed and ChIP was carried out using either anti-Six1 antibody or normal rabbit IgG. Results represent qPCR quantification of IP enrichment (expressed as percentage of input DNA) at the 48kb binding site upstream of the Slc16a10/MCT10 locus. Enrichment at the Myog proximal promoter is shown as positive control while lack of enrichment at the Hoxd10 proximal promoter is shown as specificity control. Results from each biological replicate are represented by different colors. Two-tailed paired T-test were conducted to compare enrichment with IgG and with anti-Six1. One-tailed paired T-tests also indicate that the degree of enrichment with anti-Six1 is significantly higher (p < 0.05) at the Myog and Slc16a10 loci compared to the Hoxd10 locus.

The microarray transcriptome analysis by Sakakibara et al. (Fig. 1) were acquired in mice where *Six1* was ablated around embryonic day 9, when the HSA-Cre transgene is first detected in the myotome (28,75). To explore the effect of Six1 loss-of-function specifically in adult muscles, we employed intra-muscular siRNA duplex electroporation to knockdown Six1 expression in the TA muscle (another predominantly fast-twitch muscle from muscleDB cluster 1). The TA was chosen for its small size, compared to the gastrocnemius, making electroporation more efficient. Total proteins and RNA from electroporated tissues were harvested. A western blot was performed on siRNA-treated muscle extracts demonstrating a corresponding 88% reduction of Six1 protein levels (Fig. 5A and B). The effect on target gene expression was evaluated using qRT-PCR. *Six1* mRNA expression levels decreased by approximately 57% under siSix1 knockdown condition; no compensatory effect was seen with *Six4*, a related family member (Fig. 5C). A significant decrease in MCT10 expression was observed in the *Six1* knock-down samples, validating that the expression of this TH transporter depends on *Six1* (Fig. 5C). The effects are specific to MCT10, as MCT8 expression was invariant with *Six1* knockdown. Together, these results indicate that TH transporter MCT10 is a direct target gene of *Six1* and depends on this TF to reach its normal expression level in fast-twitch skeletal muscle. It will be interesting to determine if other tissue types also depend on Six1 for MCT10 expression. In that context, it is worth noting that thyroid gland development is defective in Eya1 knockout mice (76) and that the Six1 and MCT10 mRNAs are both detected in the human adult thyroid (77).

**Figure 5.**
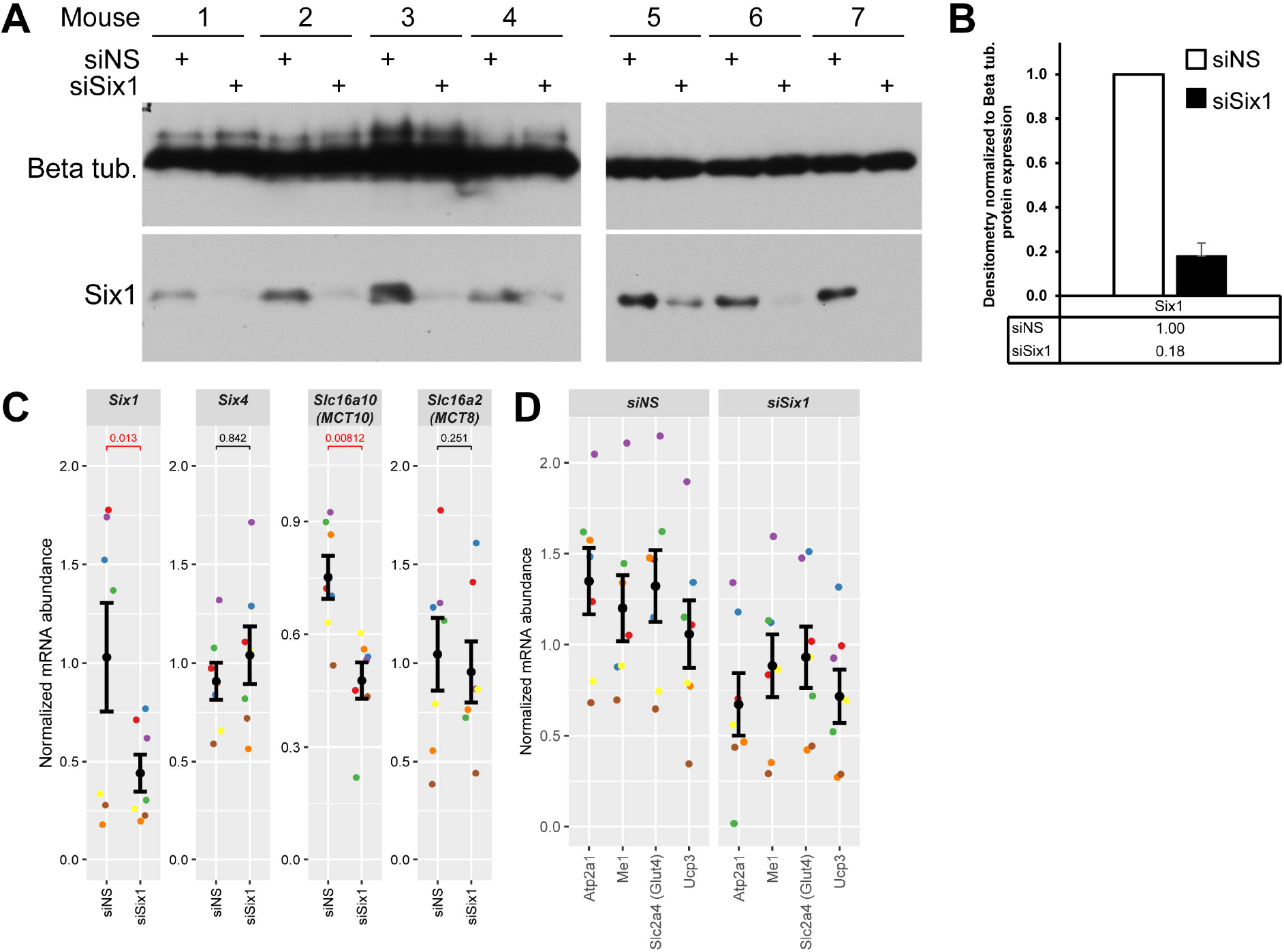
Six1 is needed for MCT10 expression and activity of the TH pathway. **A)** Western blots assaying Six1 protein expression levels under siRNA knockdown condition. siNS represents a control, non-silencing RNA duplex. Note that animals 1 to 4 and 5 to 7 were processed on different days and the western blots were done on different gels and membranes. **B)** Densitometric quantitation of Six1 abundance shown in panel A. Error bars indicate the SEM. The difference in Six1 protein abundance is statistically significant by a one-tailed paired T-Test relative to siNS (p-value < 0.05). The experiment was performed with seven mice treated identically. **C)** mRNA expression levels of *Six1* and MCT10 in electroporated mouse TA muscle after Six1 knockdown, quantified by qRT-PCR. *Six4* and MCT8 are shown as members of the same gene families with invariant expression in this experiment. mRNA levels were normalized to the geometric mean of those of 18S rRNA and *Actnb*. Data represent an average of 7 biological replicates (animals). The normalized expression data is given, with each individual replicate (mouse) shown in a distinct color. Error bars indicate the mean ± SEM of the replicates. P values in red indicate statistically significant decreases in levels of expression as determined by a one-tailed paired Student’s T-Test relative to siNS (p-value < 0.05)). **D)** mRNA expression level of a panel of four TH pathway target genes in the control and Six1 knockdown samples from panel C. Two-way ANOVA (testing mRNA levels as a function of which gene was tested and which siRNA was received) indicates significant reduction of the TH signaling gene panel with Six1 knockdown (p-value = 0.001).

### Thyroid pathway regulation is impacted negatively in Six1 loss-of-function conditions

To determine the extent of the effects of Six1 knockdown and its regulation of MCT10 expression on the thyroid regulatory pathway, a panel of four TH receptor target genes was selected for qRT-PCR analysis: *Atp2a1, Slc2a4* (Glut4), *Me1*, and *Ucp3* were chosen as they are among the best characterized direct targets of TH signaling and TH receptors in skeletal muscles (40,78–83). The downregulation of MCT10 expression with *Six1* knockdown was paralleled by that of the four TH pathway genes (Fig. 5D); the gene panel as a whole was significantly less expressed with *Six1* loss-of-function by two-way ANOVA. The panel’s reduced expression suggests a decrease in overall thyroid genomic regulatory pathway activity. We note that *Atp2a1* is a known direct target of *Six1* (17,63), and the analysis of ChIP-seq and gene expression profiling data in Fig. 1 suggests that *Me1* represents another direct target of this TF. Whether the downregulation of these genes after *Six1* loss-of-function is strictly due to decreased *Six1* activity or a combination of direct and indirect effects (*e*.*g*., through deregulation of TH signaling) remains to be formally established, but these observations raise the possibility of combinatorial regulation of gene expression by *Six1* and TH receptors. Our results also show that continued *Six1* expression is required to maintain proper gene expression programs in adult muscle.

We further examined the link between *Six1* function and TH signaling in skeletal muscle by performing gene set enrichment analysis (GSEA) on the Six1-cKO gene expression profiling data of Sakakibara *et al*. We generated two gene sets from the study of Nicolaisen et al. (84), where the impact of T3 treatment and muscle-specific knockout of the thyroid hormone receptor alpha gene (Thra-cKO) was examined by RNA-seq. Specifically, we generated a first gene set representing genes whose expression is significantly lower in T3-treated muscle of Thra-cKO compared to wild-types, and a second one with the genes showing the opposite behavior, with expression higher in the Thra-cKO tissue. We found that *Six1* activity is directly correlated with that of Thra because genes whose expression is lower in Six1-cKO are enriched for genes whose expression is lower in Thra-cKO, and vice-versa (Fig. 6). Together, these results indicate that *Six1* is functionally and positively associated with TH pathway activity in skeletal muscle.

**Figure 6.**
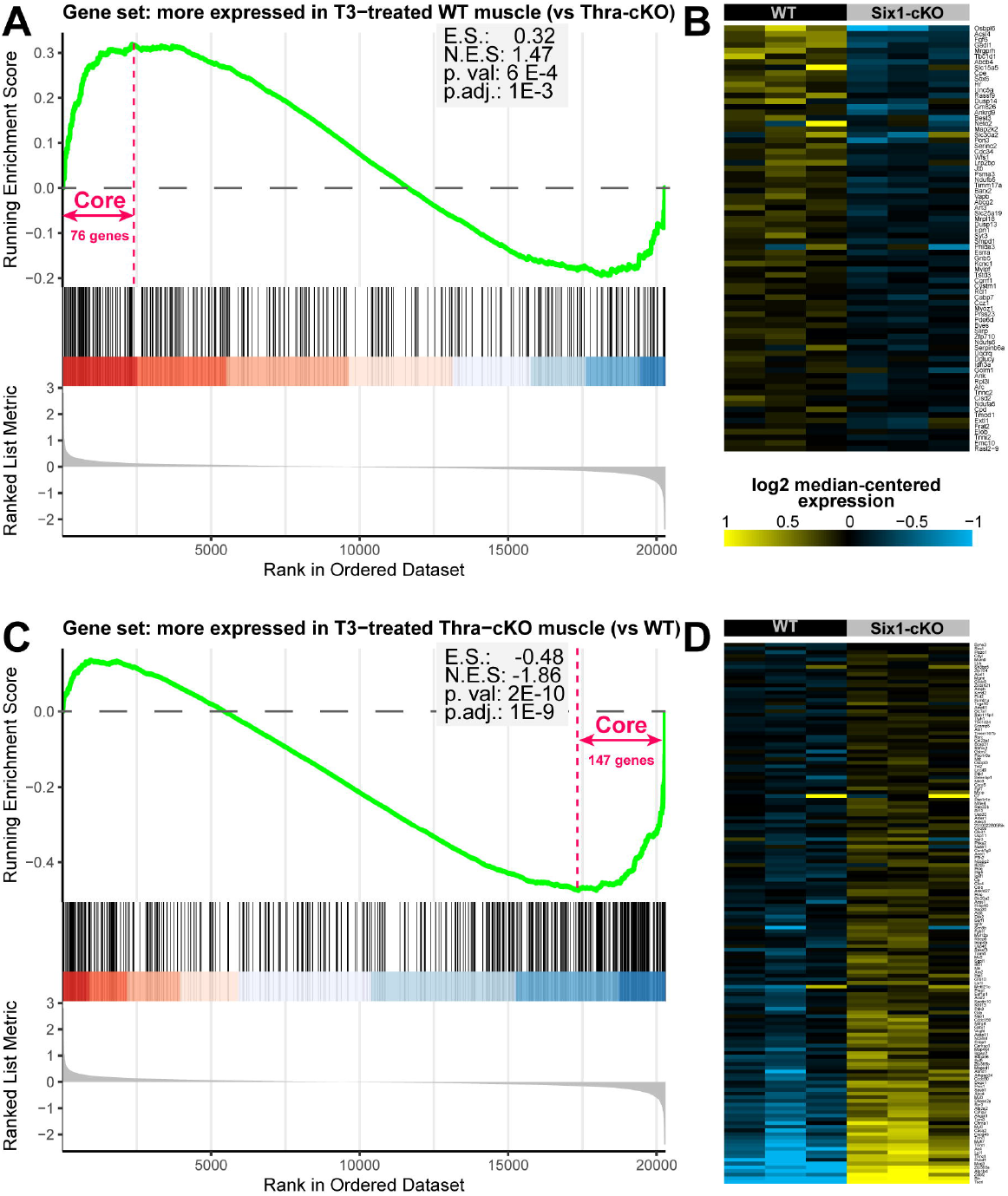
Six1 function is correlated with that of the thyroid hormone receptor transcription factor. Gene set enrichment analysis performed on the gene expression profiling of wild-type or Six1-cKO skeletal muscle (Sakakibara *et al*.), using two custom sets representing genes that are significantly less or more expressed in Thra knock-out skeletal muscle treated with T3 compared to wild-type T3-treated muscle (from Nicolaisen *et al*.). **A)** Enrichment score graph showing that when genes are ranked from the most down-regulated in Six1-cKO to the most up-regulated, the beginning of the list is enriched in genes that are more expressed in T3-treated wild-type muscle compared to Thra-cKO muscle. **B)** Heat map of the genes that contribute the most to the enrichment shown in panel A (“core enrichment” genes). The gene order from top to bottom in the heatmap follows the order from left to right in panel A. **C** and **D)** Similar analyses as in A and B but for the set representing genes that are more expressed in Thra-cKO compared to WT. The gene symbols shown in panels B and D are listed in supplementary files Code_S3 and Code_S4.

### MCT10 is required for activity of the TH signaling pathway in skeletal muscle

To examine the contribution of MCT10 down-regulation to the TH signaling phenotype observed after *Six1* loss-of-function, we used siRNA to knock down the MCT10 transporter directly. Similar to the approach taken with *Six1* knockdown, siRNA duplexes targeting the MCT10 transcript (siMCT10) were electroporated into adult TA muscle and mRNA expression levels were assessed using qRT-PCR. siMCT10 electroporation reduced MCT10 mRNA expression levels significantly and had no notable effect on the related MCT8 gene (Fig. 7A). Despite modest downregulation of MCT10 expression after siRNA knockdown (29% reduction) when compared with the efficiency of *Six1* knockdown (Fig. 5C), MCT10 knockdown was accompanied by a reduced expression of the TH signaling pathway gene panel (Fig. 7B), with trends similar to those observed under Six1 knockdown conditions (statistical significance as a group by two-way ANOVA, Fig. 5D). Finally, to further confirm that MCT10 knockdown is associated to a decrease in transcriptional response to TH, we assayed the activity of a TH-responsive luciferase reporter gene electroporated in TA muscles at the same time as the siRNA duplexes. MCT10 knockdown caused a significant reduction in the activity of the TH-dependent reporter gene (Fig. 7C). These results indicate that MCT10 is required for normal TH signaling in skeletal muscle.

**Figure 7.**
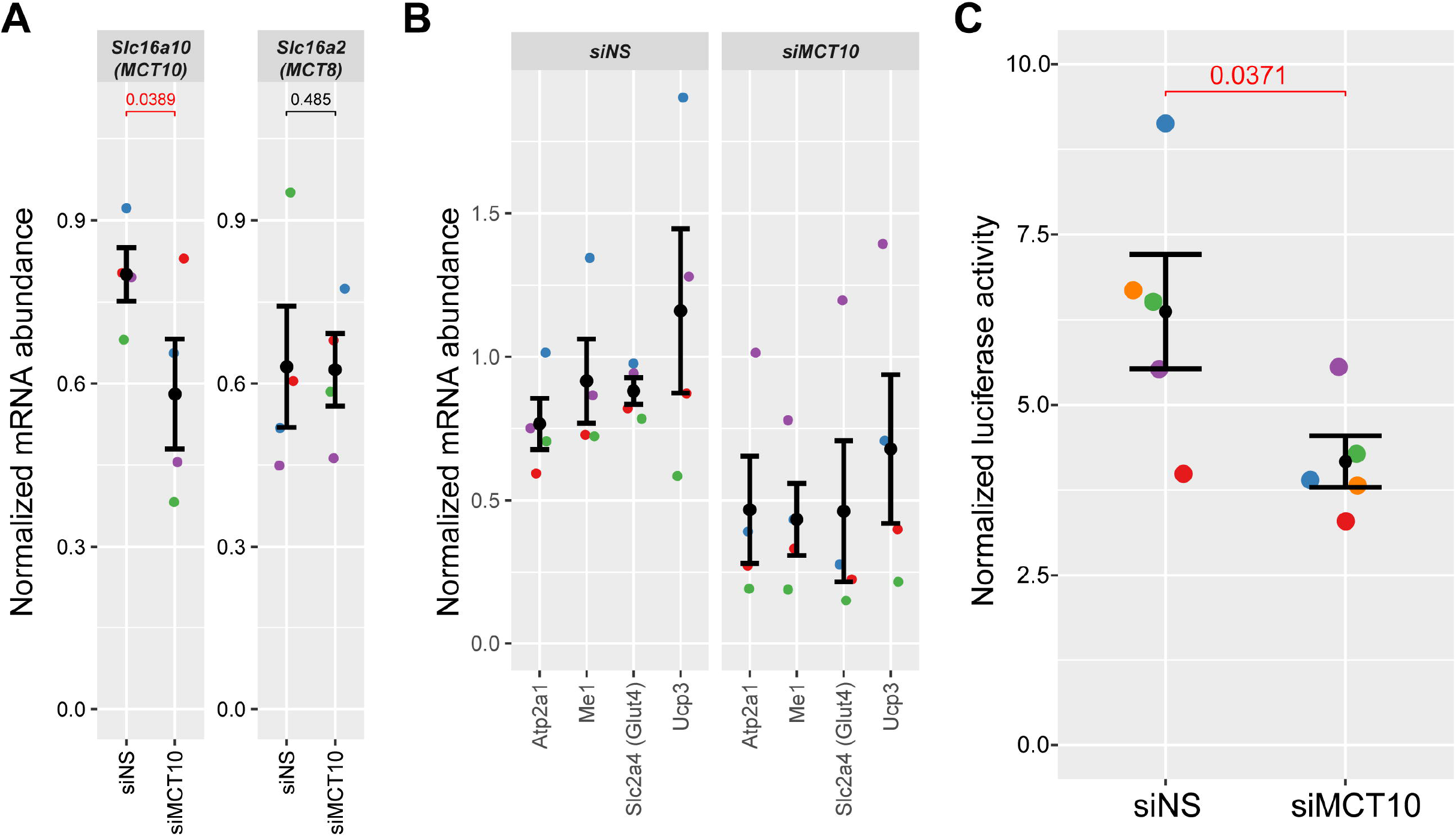
Recapitulation of TH pathway effects with MCT10 knock-down. **A)** Gene expression in TA muscles electroporated with siRNA against MCT10 or a non-silencing RNA duplex. Expression levels of MCT10, targeted by the siRNA employed, and those of the related gene MCT8 and Six1 are shown. mRNA levels are normalized to the geometric mean of 18S and Actnb. Data were obtained from 4 biological replicates (animals). Error bars indicate the mean ± SEM of the replicates. P values in red indicate statistically significant changes in levels of expression as determined by one-tailed paired Student’s T-Tests relative to siNS (p-value < 0.05). **B)** mRNA expression level of a panel of four TH pathway target genes in the control and MCT10 knockdown samples from panel A. Two-way ANOVA (testing mRNA levels as a function of which gene was tested and which siRNA was received) indicates significant reduction of the TH signaling gene panel with MCT10 knockdown (p-value = 0.005). **C)** MCT10 knock-down leads to a decrease in TH-dependent gene transcription. Relative TRE-dependent luciferase activity in TA muscle electroporated with siRNA targeting MCT10. N=5 biological replicates. The decrease in reporter gene expression is significant as determined by paired one-tailed T test, compared to non-silencing control (siNS).

As a *Six1* target gene, MCT10 is thus identified as a mediator of the influence of *Six1* on the TH signaling pathway in skeletal muscle. The implication of other transporters in TH uptake in skeletal muscle fibers cannot be ruled out. However, MCT10 appears to play a significant role in the process and other transporters are unable to fully compensate the effect of MCT10 knockdown. The situation in mature muscle appears different from that prevailing during regeneration, where MCT8 and OATP1C1 play a more important role (57). Considering the importance of TH signaling for muscle function and metabolism, the findings presented here therefore suggest that TH transporter expression must be tightly controlled, and we have shown that Six1 plays an important part in the regulatory mechanism.

## Methods

### Animal Care

Surgical procedures were performed in compliance with the University of Ottawa Animal Care and Use Committee, the Guidelines of the Canadian Council on Animal Care, and the Animals for Research Act. Female C57BL/6J mice 6–8-week-old were purchased from Charles River. Mice were sacrificed using cervical dislocation.

### ChIP-sequencing

ChIP-seq was performed for Six1 in primary mouse myoblasts as previously described (23). Sequencing data have been deposited on the NCBI Gene Expression Omnibus (GEO) under accession GSE175999. Primary myoblasts were isolated from eight weeks old female mice and were harvested sub-confluent or grown to confluence and cultured for 48 hours in differentiation medium (DMEM supplemented with 10% horse serum) to generate primary myotubes. To identify Six1 binding sites, MACS v1.4 was used with default parameters (85). Peaks called by MACS were further refined by splitting large, compound peaks into individual ones using PeakSplitter with a cut-off of 50 reads (86). Six1 ChIP-seq peaks were annotated to mouse genes using the packages *TxDb*.*Mmusculus*.*UCSC*.*mm9*.*knownGene* and *ChIPseeker* (87), setting a distance cut-off of 50 kb. Histone marks ChIP-seq and ATAC-seq in quadriceps and soleus were from the study by Barish et al. (72) and were obtained from the NCBI Short Reads Archive (accessions SRP199043 and SRP173476). To simplify the analysis, all available biological and technical replicates for each condition were combined into a single file. Reads were trimmed of adapters and low-quality sections using fastp (88), they were aligned to the mouse mm9 genome using STAR (89) in local mode with maximum intron size set to 1 base pair and using a genome index built without gene model GTF file. Only reads aligning to a single genomic location were retained. Duplicates were marked using Picard (90). Genome-wide sequencing coverage was calculated using deepTools bamCoverage, excluding reads marked as duplicates (91). In the specific case of ATAC-seq, bamCoverage was restricted to paired-end fragments between 10 and 130 base pairs, assumed to represent nucleosome-free regions. For histone mark ChIP-seq, available as single-end data, bamCoverage was performed using a read extension length calculated using SSP (92) as the most prevalent chromatin fragment size in the respective library. Genomic coverage plots were prepared using R/Bioconductor (93) and the *karyoploteR* package (94).

### Gene expression profiling data analysis

Gene expression data from the Sakakibara study (28) was retrieved from the NCBI GEO (accession GSE46151). Only gastrocnemius muscle samples were included in the analysis. Data were analyzed using R/Bioconductor and the *oligo* and *limma* packages (95,96), using RMA normalization and a differential expression testing design based on genotype. Log_2_ fold-changes were calculated, and P values were adjusted using the Benjamini-Hochberg algorithm; for significance thresholding, cut-offs of 0.58 and 0.05 were used, respectively. The list of differentially expressed genes is provided in supplementary table S1. Gene expression data from the Terry et al. muscleDB study (65) was retrieved from the NCBI GEO (accession GSE100505) and SRA accession SRP110541. Reads were trimmed with fastp and aligned to the mm10 mouse genome using STAR in local mode and a genome index built using a gene model GTF file (Mus_musculus.GRCm38.102.gtf obtained from ENSEMBL). Only reads aligning to a single genomic location were retained. Duplicates were marked, but retained for all downstream analyses, using Picard. Read summarization to genes was performed using Subread featureCounts (97). R/Bioconductor and the *DESeq2* package were used to quantitate and normalize gene expression and obtain variance stabilization expression estimates (98). Data clustering and heatmap generation were accomplished using the *pheatmap* package (99). The R script (including package versions and session information) used to analyze data and generate the figures is provided as supplementary files Code_S1. Gene set enrichment analysis was performed using the Sakakibara data and two gene sets from the Nicolaisen et al. study (84). Briefly, RNA-seq data (read counts over genes) were downloaded from NCBI GEO accession GSE146336 and analyzed in R/Bioconductor using the *edgeR* package (100). Data were normalized using TMM and batch effects were modeled using *RUVseq* with the RUVr algorithm and k=2 (101). Genes significantly up- or down-regulated in T3-treated Thra-cKO muscle compared to T3-treated wild-type muscle were identified (FDR < 0.05 with the glmQLFTest function) and saved as two separate gene sets. GSEA was performed in R/Bioconductor with the *clusterProfiler* package (102), using the fgsea algorithm with default parameters, except for the maximum gene set size limit which was adjusted to 1000. The two gene sets (with Entrez Gene ID as identifiers) are provided in supplementary file Code_S2.

### Nucleic Acid Electroporation

Electroporation experiments were performed in the tibialis anterior (TA) muscle on 6–8-week-old female C57BL/6 mice. For knockdown experiments, on day 0 of the experiment, the inter-connective tissue of the TA was digested with an intramuscular injection of 25 µL hyaluronidase (Worthington) at 0.4 unit/µL concentration through the skin of the hindlimb just above the ankle tendon, in order to ensure genetic material can surround and successfully transfect a majority of muscle fibers once injected (103). One hour later, an intramuscular injection of 22.5 µL siRNA at 20 µM was administered through the skin of the hindlimb into the belly of the TA. Electroporation of nucleic acids was performed with Tweezertrodes (2-paddle electrode assemblies, BTX-Harvard Apparatus 45-0165), used with a 7 mm gap between the electrodes. After nucleic acid material injection, ultrasound gel was applied to the skin, and two electroporation rounds were applied superficially, with the paddles in the first electroporation oriented transverse and sagittal to the hindlimb, and the second electroporation oriented transverse and contra-sagittal to the hindlimb. Electrical stimulation was carried out using a BTX ECM 630 Electroporation system programmed at a setting of 50 Volts, 6 Pulses, 50ms/pulse, and 200ms interval between pulses. Tweezertrodes electroporations were repeated on day 3 of the experiment. Non-silencing control and *Six1*- or *Mct10*-targeting siRNA duplexes were obtained from Invitrogen (Stealth siRNA technology) and reconstituted at 20 µM. The sequences are siSix1: GCGAGGAGACCAGCUACUGCUUUAA; siMct10: GCGUCUUCACAAUCCUGCUCCCUUU. The siNS negative control was Stealth RNAi siRNA Negative Control, Med GC sequence 1 (catalog number 12935300). To reduce the effect of biological variability, in all experiments, one leg received the siNS duplex while the other leg of the same animal received the Six1- or Mct10-targeting siRNA. All mice were sacrificed on day 7 of the experiment. For experiments that included a transcription reporter plasmid, the siRNA was combined with 7.5 µL of the pdV-L1 plasmid at 3 µg/µL in half-saline solution. This plasmid contains two T3 response elements and the SERCA1 basal promoter upstream of the firefly luciferase gene as well as the control renilla luciferase gene preceded by the SERCA1 promoter (kindly provided by Dr. W.S. Simonides (VU Medical Center, Amsterdam, NL) (104,105).

### Luciferase assay

Muscles were pulverized in liquid nitrogen using a chilled mortar and pestle (Plattner’s) and re-suspended in passive lysis buffer (Promega). Luciferase results were obtained by reading 5 µL of sample with 50 µL luciferase assay reagents in a Glomax Luminometer according to manufacturer’s specifications.

### RNA Extraction

All samples were extracted in 1 mL of TRIzol Reagent (Invitrogen). Dissected and pulverized TA samples were placed in TRIzol in Lysing Matrix D Tubes (MP Biomedical) and were homogenized in a MagNa Lyser machine (Roche) programmed to 7,000 RPM for a total of three 20-second bursts separated by 10-second cool-downs on ice. The samples were then spun down for 5 minutes at 12,000 g at 4ºC to remove fat content. RNA extraction proceeded as recommended by the TRIzol manufacturer. RNA was further purified by treatment with DNase I (RNase-free, New England Biolabs) and heat inactivation of the enzyme as recommended by the manufacturer.

### Reverse-Transcription

Reverse-transcription of RNA into cDNA was performed using 500 ng of DNAse-treated total RNA with the SuperScript III First-Strand Synthesis System (Thermo Fisher) using the random hexamers method and post-reaction RNAse H treatment, according to the manufacturer’s recommendation. cDNA samples were diluted to 50 µL with 10 mM Tris pH 8.0 prior to quantitative PCR.

### Chromatin Immunoprecipitation

For ChIP experiments, each replicate was performed with all hindlimb muscles from two mice. Tissue was dissected and homogenized on ice in hypotonic buffer (25 mM pH 7.8, 1.5 mM MgCl_2_, 10 mM KCl, 0.1% (v/v) NP-40 and protease inhibitors), using a Teflon Potter-Elvehjem homogenizer mounted on a benchtop drill. Formaldehyde was added from an 11X solution (11% formaldehyde, 0.1 M NaCl, 1.0 mM EDTA, 0.5 M EGTA, 50 mM HEPES pH 8.0) for 10 minutes at room temperature. Quenching was achieved using 0.125 M glycine for 5 minutes at room temperature. Samples were spun down at 1,000 g for 5 minutes at 4ºC, resuspended in fresh hypotonic buffer, filtered through 70 µm cell strainer, spun down again at 1,000 g for 5 minutes at 4ºC and re-suspended in 650 µL sonication buffer (10 mM EDTA, 50 mM Tris-HCl pH 8, 0.1% (w/v) SDS and protease inhibitors). The nuclear pellet was collected by centrifugation and lysed in nuclear lysis buffer (200 mM NaCl, 1 mM EDTA, 0.5 mM EGTA, 10 mM Tris-HCl pH 8 and protease inhibitors). Nuclei were pelleted and resuspended in 650 µL sonication buffer (1 mM EDTA, 0.5 mM EGTA, 0.5% (w/v) sodium sarkosyl, 10 mM Tris-HCl pH 8.0 and protease inhibitors). All sonications were carried out using a probe sonicator at an amplitude of 40% power, with 60 cycles of 1 second on to shear DNA and 4 seconds off to cool down. The rest of the assay was performed as previously published (17), using 25 µg of chromatin per sample, pre-clearing with Protein A sepharose beads, using 2 µg of antibody per sample with overnight incubation, seven washes of the beads, elution in SDS combined with cross-link reversal and DNA clean-up by phenol extraction followed by ethanol precipitation. Primary antibodies used in ChIP were anti-Six1 (rabbit polyclonal HPA001893, Sigma) and normal rabbit IgG (Jackson ImmunoResearch). The final pellet was re-suspended in 50 µL 10 mM Tris-HCl pH 8.0 for downstream qPCR.

### qPCR

qPCR on cDNA and ChIP samples was carried out in reactions containing SYBR Green I HotStarTaq enzyme (Qiagen). Relative quantitation was performed using standard curves made by pooling aliquots of each cDNA sample being examined. Since absolute mRNA abundance of each gene varies in the pool, the expression of a given gene can be compared across samples, but the expression levels of different genes cannot be directly compared. Instead, standard curves for qChIP were made with input genomic DNA; assuming every gene is present in the same allele number in the genome, enrichment levels of different genes can be compared directly. All samples were run as technical triplicates and their geometric mean was taken. For qRT-PCR, relative expression was obtained by dividing by the geometric mean of 18S ribosomal RNA and *Actnb*, which were found invariable under experimental conditions (106). For each experiment, the graphs show individual biological replicates (experimental mice) as separate points, along with their mean and standard errors. Oligonucleotide sequences are provided in Table S2.

### Protein Extraction, Quantitation and Immunodetection

Muscle proteins were extracted from the TRIzol organic phase following the procedure recommended by the manufacturer. Protein pellets were resolubilized in Urea-SDS Sample Buffer (6 M Urea, 1 % SDS, 20 mM Tris pH 6.8) at a ratio of about 3 µL buffer per 1mg of tissue and were allowed to solubilize with gentle rocking overnight at 4ºC. Sample protein concentrations were determined using the BCA Protein Assay Kit (Thermo Scientific). Primary antibodies were anti-Six1 (rabbit polyclonal HPA001893, Sigma) and anti-beta-Tubulin (mouse monoclonal hybridoma clone E7, DSHB). Secondary antibodies conjugated with HRP were used. Signal was acquired on X-ray films, which were subsequently digitized and analyzed in the FIJI software for densitometric quantitation.

### Statistical Analyses

Required number of replicates was determined empirically for sufficient statistical power. No samples were excluded specifically from analysis. R and the *rstatix* package were used to carry out statistical analyses (107). Statistical significance for qPCR, densitometry, and luciferase assays were determined by paired T-tests, always pairing the experimental muscle electroporated with knockdown siRNA, with its contralateral muscle from the same animal electroporated with siNS. In the presence of prior data suggesting the direction of the effects, one-tailed tests were used. P-Values ≤ 0.05 were considered to represent means that were significantly different. Where applicable, multiple hypothesis adjustment of p-values was performed using the Benjamini-Hochberg method(108). Unless otherwise stated, no fewer than 3 biological or technical replicates were obtained per experiment. Statistically analyzed control sample groups and experimental sample groups always utilized the same number of technical and biological replicates. In the case of mRNA expression of the TH signaling gene panel (composed of *Atp2a1, Me1, Slc2a4* and *Ucp3*) after *Six1* or MCT10 knockdown, a two-way ANOVA test of the effect of independent categorical variables “Gene tested” and “siRNA duplex used” on the dependent continuous variable “Normalized expression” was performed. Normal distribution and variance homogeneity assumptions were confirmed by Shapiro-Wilk’s and Levene’s tests, respectively. P-values for the effects of “siRNA duplex used” were significant and are reported.

## Acknowledgements

The authors would like to thank the following individuals or groups for their contributions: Sarah Hemens and Jack Guthrie for experimental assistance; the staff of the animal care and veterinary services of the University of Ottawa for technical assistance; the McGill University and Génome Québec Innovation Centre for ChIP-seq library preparation and high-throughput sequencing; Compute Canada and technical staff for high-performance computing access and support; and Warner Simonides (Amsterdam, The Netherlands) for the TRE reporter plasmid. The E7 hybridoma against beta-tubulin, developed by M. Klymkowsky was obtained from the Developmental Studies Hybridoma Bank, created by the NICHD of the NIH and maintained at The University of Iowa, Department of Biology, Iowa City, IA 52242. This work was supported by a Canadian Institutes of Health Research operating grant (MOP 119458) to A. B..

